## Supplementary Materials for "PDGFRα signaling regulates Srsf3 transcript binding to affect PI3K signaling and endosomal trafficking"

Thomas E. Forman *et al*.

**The PDF file includes:**

Figs. S1 to S6
Tables S1, S7 and S16

**Other Supplementary Material for this manuscript includes the following**:

Tables S2-S6 and S8-S15

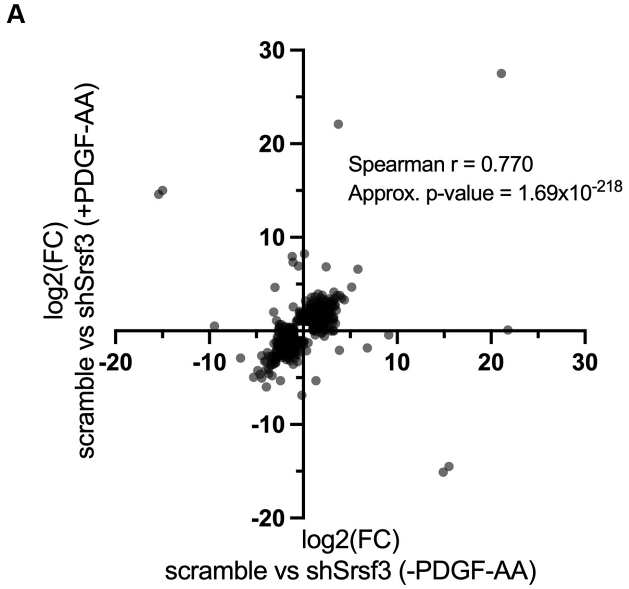


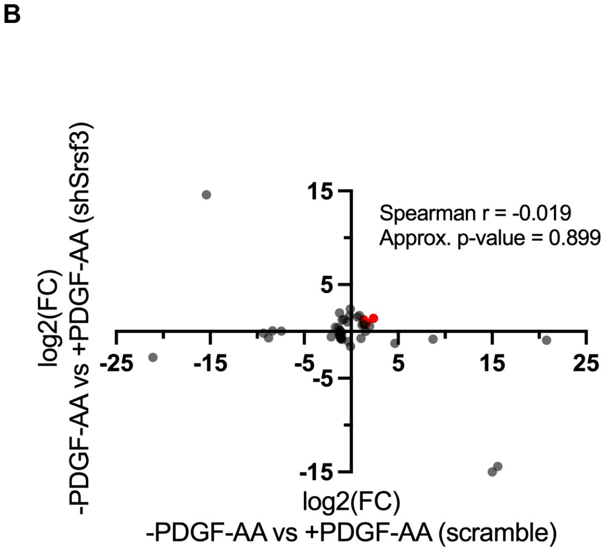


**Figure S1. High correlation of Srsf3-dependent differentially-expressed genes across ligand treatment conditions. (A,B)** Scatter dot plots depicting Srsf3-dependent (A) and PDGF-AA-dependent (B) differentially-expressed genes. $Log_2$(fold change) (FC) values represent $log_2$(shSrsf3 normalized counts/scramble normalized counts) (A) or $log_2$(+PDGF-AA normalized counts/-PDGF-AA normalized counts) (B). Spearman correlation values and approximate p-values are listed. Immediate early genes are represented in red in B.

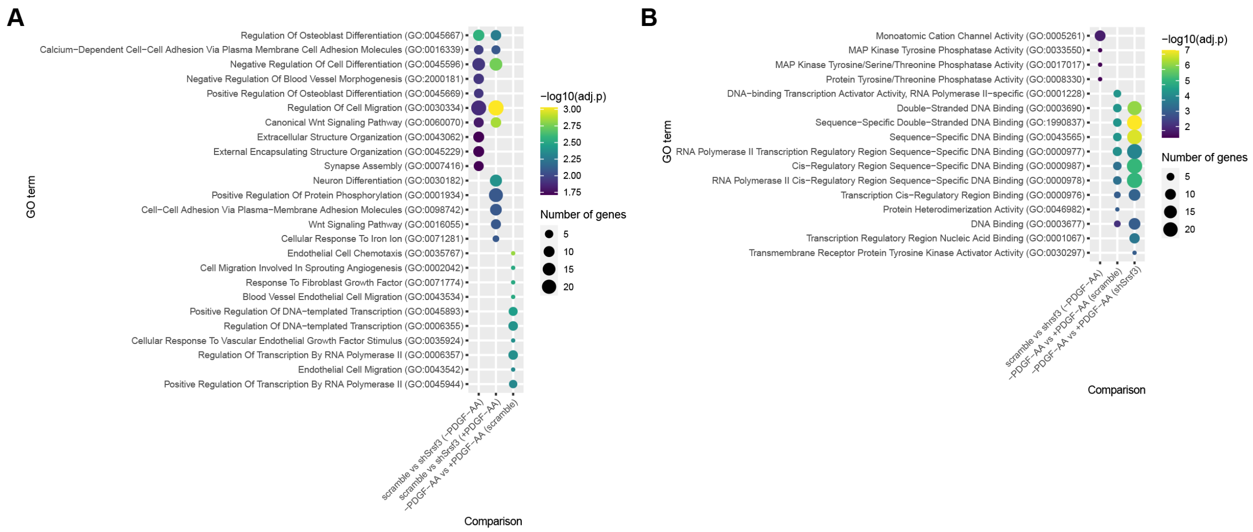


**Figure S2. Gene ontology analysis of differentially-expressed genes across treatment comparisons. (A,B)** Bubble plots depicting up to ten of the most significant gene ontology (GO) terms for biological process (A) and molecular function (B) for Srsf3-dependent and PDGF-AA-dependent differentially-expressed genes. Colors correspond to -log10(adjusted p-value); sizes correspond to number of genes.

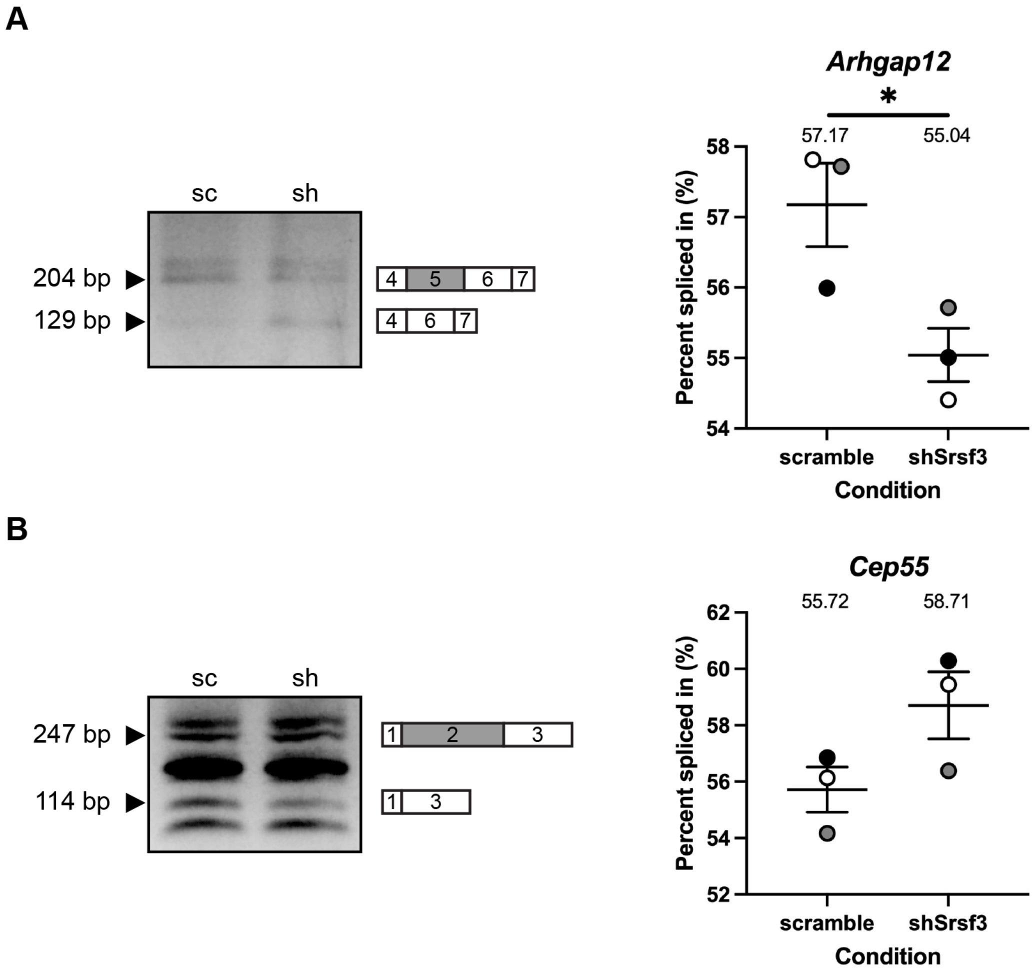


**Figure S3**. **qPCR validation of differential AS between scramble and shSrsf3 samples.** **(A,B)** Representative qPCR gels (left) with depictions of differentially alternatively-spliced exon (gray), and upstream and downstream sequences (white) that were assessed by qPCR in scramble (sc) versus shSrsf3 (sh) samples for *Arhgap12* (A) and *Cep55* (B). Scatter dot plots (right) depicting the percent spliced in from $n$ = 3 biological replicates as at left. Data are mean ± s.e.m. *, $P < 0.05$. Shaded circles correspond to independent experiments.

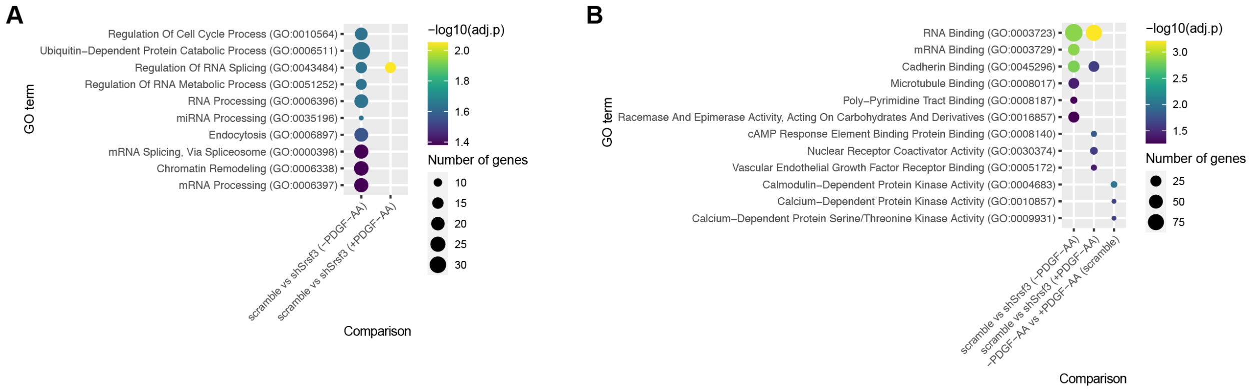


**Figure S4. Gene ontology analysis of alternatively-spliced transcripts across treatment comparisons. (A,B)** Bubble plots depicting up to ten of the most significant gene ontology (GO) terms for biological process (A) and molecular function (B) for Srsf3-dependent and PDGF-AA-dependent alternatively-spliced transcripts. Colors correspond to -log10(adjusted p-value); sizes correspond to number of genes.

| -PDGF-AA | | | | +PDGF-AA | | | |
|---|---|---|---|---|---|---|---|
| **Sequence logo** | **Occurrence in eCLIP (per 1000 peaks)** | **Occurrence in control (per 1000 peaks)** | **p-value** | **Sequence logo** | **Occurrence in eCLIP (per 1000 peaks)** | **Occurrence in control (per 1000 peaks)** | **p-value** |
| 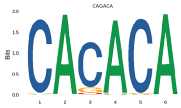 | **4095** | **991** | **$<2.2 \times 10^{-16}$** | 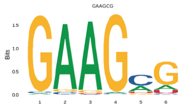 | **1432** | **465** | **$<2.2 \times 10^{-16}$** |
| 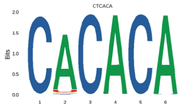 | **4281** | **1209** | **$<2.2 \times 10^{-16}$** | 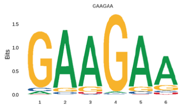 | **2032** | **649** | **$<2.2 \times 10^{-16}$** |
| 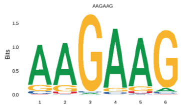 | **1307** | **555** | **$3.6 \times 10^{-11}$** | 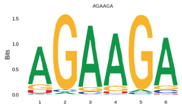 | **2029** | **726** | **$<2.2 \times 10^{-16}$** |
| 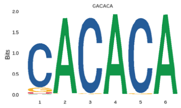 | **4239** | **1135** | **$<2.2 \times 10^{-16}$** | 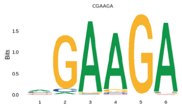 | **1279** | **353** | **$<2.2 \times 10^{-16}$** |
| 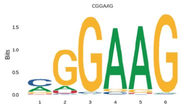 | **602** | **253** | **$5.4 \times 10^{-15}$** | 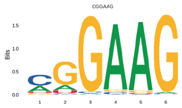 | **833** | **315** | **$<2.2 \times 10^{-16}$** |
| 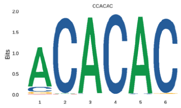 | **4286** | **1151** | **$<2.2 \times 10^{-16}$** | 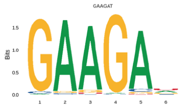 | **1273** | **436** | **$<2.2 \times 10^{-16}$** |
| 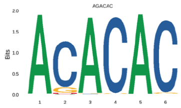 | **4169** | **1116** | **$<2.2 \times 10^{-16}$** | 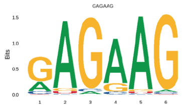 | **1660** | **663** | **$<2.2 \times 10^{-16}$** |
| 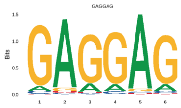 | **1233** | **488** | **$<2.2 \times 10^{-16}$** | 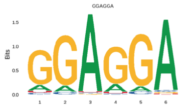 | **1486** | **636** | **$7.5 \times 10^{-16}$** |
| 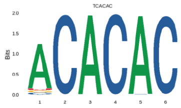 | **3970** | **1026** | **$<2.2 \times 10^{-16}$** | 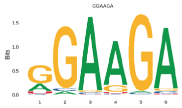 | **1726** | **631** | **$<2.2 \times 10^{-16}$** |
| 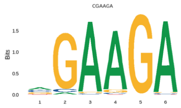 | **758** | **307** | **$3.0 \times 10^{-10}$** | 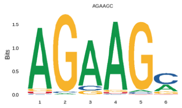 | **1383** | **545** | **$<2.2 \times 10^{-16}$** |

**Figure S5. PDGFRα signaling influences Srsf3 binding specificity.** Top 10 motifs enriched in eCLIP peaks in the absence (left) or presence (right) of PDGF-AA stimulation with associated *P* values.

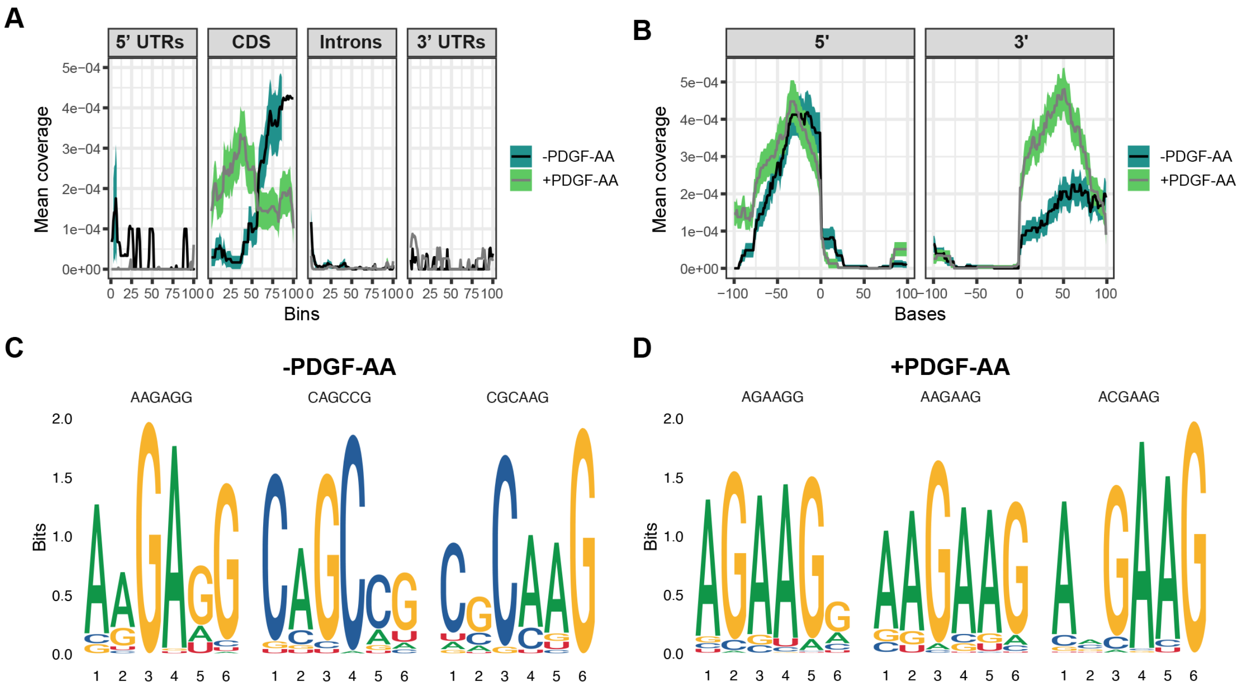


**Figure S6. Srsf3 exhibits differential transcript binding upon PDGFRα signaling in the subset of transcripts from the high-confidence, overlapping datasets. (A,B)** Mean coverage of eCLIP peaks within the high-confidence, overlapping datasets across various transcript locations (A) and surrounding the 5' and 3' splice sites (B) in the absence or presence of PDGF-AA stimulation. **(C,D)** Top three motifs enriched in eCLIP peaks within the high-confidence, overlapping datasets in the absence (C) or presence (D) of PDGF-AA stimulation.

**Table S1. RNA-seq sample information.**

| Sample | Raw read pairs | Trimmed read pairs for Salmon input | Salmon mapping rate | Trimmed read pairs (125 bp) for STAR input | STAR unique mapping rate |
|---|---|---|---|---|---|
| -PDGF-AA scramble_1 | 47181591 | 44410442 | 0.89055 | 36343779 | 0.8773 |
| -PDGF-AA scramble_2 | 54612492 | 50500367 | 0.878847 | 39971864 | 0.8681 |
| -PDGF-AA scramble_3 | 69353787 | 65529075 | 0.912399 | 48327896 | 0.9022 |
| -PDGF-AA shSrsf3_1 | 91657568 | 84086217 | 0.913324 | 61269254 | 0.9035 |
| -PDGF-AA shSrsf3_2 | 77309551 | 71220292 | 0.91638 | 49390634 | 0.9013 |
| -PDGF-AA shSrsf3_3 | 42645900 | 41078549 | 0.910338 | 28737018 | 0.9054 |
| +PDGF-AA scramble_1 | 71080979 | 66836059 | 0.916828 | 48116755 | 0.9027 |
| +PDGF-AA scramble_2 | 69667521 | 64974624 | 0.890505 | 47762451 | 0.884 |
| +PDGF-AA scramble_3 | 78680108 | 72721916 | 0.911689 | 52936280 | 0.9008 |
| +PDGF-AA shSrsf3_1 | 42776470 | 41165756 | 0.914797 | 28076373 | 0.9019 |
| +PDGF-AA shSrsf3_2 | 37944773 | 35759828 | 0.908637 | 23528077 | 0.8987 |
| +PDGF-AA shSrsf3_3 | 36391090 | 34257983 | 0.911463 | 26455873 | 0.8995 |

**Table S2. DEseq2 output.**

**Table S3. rMATS output for scramble (-PDGF-AA) versus shSrsf3 (-PDGF-AA) RNA-seq analysis.**

**Table S4. rMATS output for scramble (+PDGF-AA) versus shSrsf3 (+PDGF-AA) RNA-seq analysis.**

**Table S5. rMATS output for -PDGF-AA (scramble) versus +PDGF-AA (scramble) RNA-seq analysis.**

**Table S6. rMATS output for -PDGF-AA (shSrsf3) versus +PDGF-AA (shSrsf3) RNA-seq analysis.**

**Table S7. eCLIP sample information.**

| Sample | Raw read pairs | Trimmed read pairs | Collapsed reads | Reads after removing repetitive elements | Mapped reads | Peaks | Annotated Peaks |
|---|---|---|---|---|---|---|---|
| -PDGF-AA size-matched input | 34303575 | 22904092 | 13449745 | 13358235 | 17206 | 6969 | 6607 |
| -PDGF-AA replicate 1 | 22983544 | 18369023 | 2758371 | 2758371 | 440436 | | |
| -PDGF-AA replicate 2 | 15666256 | 12263540 | 2742638 | 2065674 | 388996 | | |
| +PDGF-AA size-matched input | 52420337 | 37454316 | 13811675 | 13643948 | 24275 | 9075 | 8623 |
| +PDGF-AA | 30105052 | 23355466 | 3417325 | 2845801 | 872085 | | |

**Table S8. eCLIP output.**

**Table S9. Raw peak counts of eCLIP peaks across various transcript locations.**

**Table S10. Matt output.**

**Table S11. List of transcripts and genes from Venn diagram in Figure 5A.**

**Table S12. High confidence, overlapping dataset output correlating eCLIP with scramble (-PDGF-AA) versus shSrsf3 (-PDGF-AA) rMATS RNA-seq analysis.**

**Table S13. High confidence, overlapping dataset output correlating eCLIP with scramble (+PDGF-AA) versus shSrsf3 (+PDGF-AA) rMATS RNA-seq analysis.**

**Table S14. High confidence, overlapping dataset output correlating eCLIP with -PDGF-AA (scramble) versus +PDGF-AA (scramble) rMATS RNA-seq analysis.**

**Table S15. High confidence, overlapping dataset output correlating eCLIP with -PDGF-AA (shSrsf3) versus +PDGF-AA (shSrsf3) rMATS RNA-seq analysis.**

**Table S16. Primers used in qPCR analysis.**

| Transcript | Forward primer (5' to 3') | Reverse primer (5' to 3') |
|---|---|---|
| *Arhgap12* | GGAGACATAGCACCATTGTG | GCACTGCCCAAGAAGACAAC |
| *Cep55* | CCTTTCGGCTCCTTTGAACT | GCAGTGTCTGACTTGGAGCT |
| *Wdr81* | GCTTTGTGGACTGCAGGAAG | GCAGGGAACAGACACCAATC |
